# Softer substrates mechanical primes durable and metabolically fit CD8+ T cells for anti-cancer activity

**DOI:** 10.64898/2026.04.22.720068

**Authors:** Aseel Alatoom, Jiranuwat Sapudom, Kamil Elkhoury, Muhammedin Deliorman, Mostafa Khair, Sanjairaj Vijayavenkataraman, Mohammad Qasaimeh, Jeremy Teo

**Affiliations:** Laboratory for Immuno Bioengineering Research and Applications, Division of Engineering, New York University Abu Dhabi, Abu Dhabi, UAE; The Vijay Lab, Division of Engineering, New York University Abu Dhabi, Abu Dhabi, United Arab Emirates; Advanced Microfluidics and Microdevices Laboratory, Division of Engineering, New York University Abu Dhabi, Abu Dhabi, UAE; Core Technology Platforms, New York University Abu Dhabi, Abu Dhabi P.O. Box 129188, United Arab Emirates; Department of Mechanical and Aerospace Engineering, Tandon School of Engineering, New York University, Brooklyn, NY 11201, USA; Department of Biomedical Engineering, New York University, New York, USA; Department of Chemical Engineering, American University of Sharjah, Sharjah, United Arab Emirates

**Keywords:** CD8+ T cells, mechanobiology, polyacrylamide hydrogels, adoptive cell therapy, solid tumors, T cell metabolism

## Abstract

Adoptive T cell therapy for solid tumors is limited by poor persistence of CD8^+^ T cells, a dysfunction that is often programmed during ex vivo expansion. Here, we show that, when biochemical inputs are held constant,substrate mechanics alone can direct durable anti-tumor function in primary human CD8^+^ T cells. Using polyacrylamide (PA) hydrogels of defined stiffness (soft ∼1 kPa; stiff ∼55 kPa) in both flat and bead formats, we first establish that contact geometry dominates early activation, whereas substrate stiffness governs the 14-day expansion trajectory. Across the rapid expansion protocol, flat PA substrates sustain proliferation, limit PD-1/LAG-3 acquisition, and preserve a balanced effector-regulatory cytokine profile. In contrast, Dynabeads™-expanded cells exhibit net cell loss and a more pronounced decline in cytokine output over time. To define the underlying programs, RNA-seq identifies a 125-gene biomimetic core shared by both PA conditions but absent from Dynabeads™, encompassing proliferation, OXPHOS, mechanobiology, and a stem-like precursor (Tpex) signature. Consistent with these transcriptional differences, metabolic profiling shows that flat soft PA best preserves dual glycolytic and mitochondrial capacity at day 14, indicating enhanced bioenergetic flexibility. Functionally, PA-primed CD8^+^ T cells display superior cytotoxicity against MDA-MB-231 and MCF-7 breast cancer cells in both 2D and collagen-based 3D co-cultures, with this advantage maintained under matrix constraints that mimic solid tumor microenvironments. Together, these findings establish substrate mechanics as a tunable and functionally decisive design parameter for engineering durable, solid-tumor-effective CD8^+^ T cell products in preclinical in vitro models of solid tumors.

## 1. Introduction

Adoptive cell therapy (ACT) has transformed the treatment of relapsed B-cell malignancies, with chimeric antigen receptor (CAR)-T products achieving durable complete remissions in patients refractory to standard therapy [1, 2]. In contrast, in solid tumors,which account for over 90% of cancer-related deaths,objective response rates typically remain below 30% [3]. This translational gap reflects not a failure of target recognition, but a failure of T cell persistence. Following infusion, CD8^+^ T cells must navigate dense stromal barriers, survive hypoxic and metabolically restrictive niches, and withstand coordinated immunosuppressive signals that drive progressive dysfunction [4-6]. Importantly, a substantial component of this suppression is physical. Tumor stiffening and collagen densification restrict CD8^+^ T cell motility and effector function while engaging YAP-mediated mechanosignalling pathways that further suppress activity [7, 8]. Critically, however, much of the dysfunction observed after infusion is not acquired de novo within the tumor, but is instead pre-programmed during ex vivo expansion [9-11]. T cell manufacturing therefore lies at the centre of the solid-tumor durability problem, with the activation platform acting as a key determinant of post-infusion fate. Current clinical-standard expansion relies on anti-CD3/CD28 Dynabeads™, which provide potent activation but drive rapid terminal differentiation, premature checkpoint expression, and bioenergetic collapse [12-14]. These phenotypes are closely associated with poor persistence and diminished anti-tumor efficacy, highlighting a central limitation of existing manufacturing strategies.

CD8^+^ T cells are inherently mechanosensitive, and the biophysical properties of the activation interface play a central role in shaping immune synapse (IS) formation, cytoskeletal remodeling, and downstream signalling [15-18]. In this context, substrate stiffness and geometry regulate force-dependent TCR-pMHC catch-bond engagement [19, 20], immune synapse architecture [21, 22], and actin dynamics mediated by Arp2/3 and WASP-dependent cytoskeletal tension [23-25]. These processes are further coupled to signalling through mechanosensitive channels such as PIEZO1, linking mechanical inputs directly to transcriptional outcomes [26, 27]. Early studies established that increasing substrate stiffness enhances naïve CD8^+^ T cell activation [17], and subsequent work has extended this principle to broader functional outcomes, including regulatory T cell induction [28], CD8^+^ effector differentiation under 3D confinement [18], and the generation of functionally distinct T cell populations using viscoelastic biomaterials [29]. Together, these findings establish the mechanical properties of the activation interface as a fundamental regulator of T cell fate, rather than a secondary or permissive factor.

Despite these advances, clinical-grade T cell expansion systems remain mechanically non-physiological. Anti-CD3/CD28 is typically presented on rigid polystyrene beads with elastic moduli orders of magnitude higher than the ∼1-10 kPa range of lymphoid tissue [30, 31]. While biomimetic alternatives, including lipid-bilayer-coated microparticles [32], alginate scaffolds [33], and mesoporous silica rods [9], can enhance acute activation, systematic comparisons against the clinical standard across the full 14-day rapid expansion protocol (REP) remain limited. Moreover, few studies integrate activation dynamics with transcriptional, metabolic, and functional cytotoxic readouts in physiologically relevant 3D tumor models. As a result, it remains unclear whether early activation gains translate into durable anti-tumor function.

We hypothesized that biomimetic substrate mechanics can program CD8^+^ T cells toward a metabolically resilient and functionally durable state that outperforms Dynabeads™ in clinically relevant settings. To test this, we established an integrated mechanism-to-function pipeline anchored to the 14-day REP. Polyacrylamide (PA) substrates of defined stiffness (soft ∼1 kPa; stiff ∼55 kPa) were engineered in both flat and bead formats and benchmarked directly against Dynabeads™. We first asked whether contact geometry governs early activation (day 7), and whether substrate mechanics sustain activation across the full expansion window (days 2-14). We then defined the transcriptional and metabolic programs that distinguish PA-expanded from Dynabeads™-expanded cells, and finally tested whether these differences translate into cytotoxic function against triple-negative (MDA-MB-231) and luminal (MCF-7) breast cancer targets in both 2D and matrix-constrained 3D co-culture. By linking each layer of analysis sequentially, this framework enables direct evaluation of how mechanical inputs shape durable anti-tumor immunity.

## 2. Materials and methods

### 2.1 Hydrogel preparation and functionalization

Anti-CD3/anti-CD28-functionalized polyacrylamide (PA) gels were prepared as previously described [34]. Hydrogels were polymerized on glutaraldehyde-treated 13 mm glass coverslips (VWR, Germany) by varying acrylamide/bis-acrylamide ratios to generate soft (∼1 kPa) and stiff (∼55 kPa) substrates; polymerization was initiated with ammonium persulfate/TEMED (Thermo Fisher Scientific, USA). Surfaces were functionalized with 1 µg/mL ultra-LEAF anti-human CD3 (clone OKT3) and anti-CD28 (clone CD28.2) (BioLegend, USA).

PA beads were fabricated by water-in-oil inverse emulsification in vegetable oil containing 5% (v/v) Span-85, with aqueous PA injected at 90 µL/min (New Era Pump Systems, USA) under stirring (IKA, Germany), and functionalized as above. Dynabeads™ Human T-Activator CD3/CD28 (Gibco, USA) served as the clinical-standard comparator.

### 2.2 Mechanical characterization

Macroscopic viscoelastic properties were measured by shear rheology (MCR 702, Anton Paar, Austria; 25 mm parallel plates, 1 mm gap, 1 Hz isothermal time sweep) to obtain storage modulus (G′) and complex viscosity (η), in triplicate.

Microscale Young’s modulus (E) was determined by AFM force spectroscopy (JPK Nanowizard 4 XP, Bruker, Germany) using a 6.62 µm SiOL colloidal probe (CP-PNPL-SiO-C, NanoAndMore, Germany; k ≈ 0.08 N/m), calibrated by the thermal noise method [35]. Force curves (32 × 32 grid over 10 × 10 µm; 4 µm/s approach; 3 nN setpoint) were acquired on gels in glass-bottom dishes (Ibidi, Germany) and analyzed with PyJibe using a Sneddon model [36].

### 2.3 CD8^+^ T cell isolation and expansion

Cryopreserved PBMCs from five independent donors (AllCells, USA) were thawed, and CD8^+^ T cells were isolated by negative selection (BioLegend, USA). Purity (>95%) was confirmed by BV650 anti-CD8α (clone HIT8a, BioLegend) staining on an Attune NxT flow cytometer (Thermo Fisher Scientific, USA).

Cells were seeded at 2 × 10^5^ cells/mL on soft or stiff PA gels, PA beads, or Dynabeads™ in RPMI-1640 supplemented with 10% FBS, 1% penicillin/streptomycin, HEPES, sodium pyruvate, 0.01% β-mercaptoethanol, and 100 U/mL recombinant human IL-2 (BioLegend, USA). A 3:1 bead-to-cell ratio was used for all bead conditions, consistent with standard Dynabeads™protocols [14]. Medium was replenished every 2 days. Cell numbers were determined every at days 2, 4, 7, 10, 12 and 14 using the Attune NxT flow cytometer, and population doubling level (PDL) was calculated as PDL = log_2_(N_counted_ /N_seeded_), where N_counted_ is the cell number at each time point and N_seeded_ is the initial seeding number of 2 × 10^5^ cells/mL.

### 2.4 Activation, exhaustion, and cytokine profiling

Cell surface marker staining was performed using PE/Fire 640 anti-CD69 (FN50), FITC anti-CD44 (BJ18), BV421 anti-PD-1 (EH12.2H7), and PE anti-LAG-3 (11C3C65) (all BioLegend, USA), and analyzed on an Attune NxT flow cytometer. Cytokine secretion in culture supernatants at days 4, 7, and 14 was quantified using a bead-based multiplex assay (LEGENDplex™ Human CD8/NK Panel, BioLegend, USA), measuring IL-2, IL-4, IL-6, IL-10, IL-17A, IFN-γ, TNF-α, granzymes A/B, perforin, granulysin, sFas, and sFasL. Data were analyzed using Qognit software (BioLegend, USA). All assays were performed in duplicate per donor.

### 2.5 Metabolic profiling

Oxygen consumption rate (OCR) and extracellular acidification rate (ECAR) were measured using a Seahorse XF96 analyzer (Agilent, USA) at days 4, 7, and 14 post-activation. Cells were reconditioned in XF RPMI (pH 7.4) supplemented with glucose, pyruvate, and glutamine (Agilent), seeded at 2 × 10L cells per well, and equilibrated at 37 °C without COL for 40 min.

Following baseline measurements, cells received sequential injections of 0.5 µM rotenone/antimycin A (Rot/AA) to inhibit mitochondrial respiration and 50 µM 2-deoxy-D-glucose (2-DG) to inhibit glycolysis. These measurements enabled calculation of glycolytic proton efflux rate (GlycoPER) and compensatory glycolysis. Values were normalized to cell number, determined from parallel counting wells processed identically. Experiments were performed in triplicate per donor.

### 2.6 Bulk nanopore RNA sequencing

CD8^+^ T cells expanded for 7 days on Dynabeads™, soft PA gels, or stiff PA gels were profiled in biological triplicate (three independent donors per condition, n = 3). Day 7 was selected as the time point of peak functional engagement across platforms, enabling transcriptomic comparison prior to divergence in exhaustion trajectories. Total RNA was extracted using TRIzol (Thermo Fisher Scientific, USA). RNA quantity and integrity were assessed using the Qubit RNA IQ Assay Kit, and only samples with IQ > 4 were used for library preparation. Libraries were prepared using the Oxford Nanopore Technologies (ONT) cDNA protocol and sequenced on a PromethION platform. Reads passing quality filters were aligned to GRCh38 (GENCODE v45) using minimap2 [37] in splice-aware mode, and gene-level counts were generated with featureCounts [38]. Counts were filtered, variance-stabilized, and analyzed by PCA. Ten curated gene signatures (activation [CD44], Tpex, proliferation [MYC + ribosome], mechanobiology, MHC-I, OXPHOS, glycolysis, chemokines, cytotoxicity [GZM/PRF], and TCA cycle) were scored as the mean z-scored expression of constituent genes. Statistical significance was assessed by permutation testing. Condition-specific master genes were ranked based on fold-change and specificity. A transcriptional metabolic map was constructed using OXPHOS and glycolysis scores as proxies for OCR and ECAR, respectively.

### 2.7 Tumor cell co-culture and cytotoxicity

MDA-MB-231 (triple-negative, mesenchymal-like) and MCF-7 (luminal, epithelial) breast cancer cells were cultured in high-glucose DMEM supplemented with 10% FBS and 1% penicillin/streptomycin (Gibco, USA). For 2D assays, tumor cells were seeded at 5 × 10^4^ cells/mL in 96-well plates and allowed to adhere overnight. For 3D assays, wells were treated with 15% NaOH (30 min), functionalized with 1% APTES and 0.5% glutaraldehyde, and overlaid with 2 mg/mL reconstituted collagen type I matrices [39]. Tumor cells were embedded and cultured overnight prior to T cell addition.

Activated CD8^+^ T cells (expanded for 4, 7, or 14 days) were added at a 3:1 effector-to-target ratio and co-cultured for 72 h. Cytotoxicity was assessed by flow cytometry using CellEvent™ Caspase-3/7 Green (1:200) and DRAQ7™ (1:1000). Tumor cells were gated by FSC/SSC. Killing was expressed as the percentage of dead tumor cells, with apoptotic (caspase-3/7LDRAQ7L) and necrotic (DRAQ7L) fractions distinguished accordingly.

### 2.8 Statistical analysis

Data are presented as mean ± SD from five donors with duplicate measurements per condition unless otherwise specified. Two-group comparisons were performed using the Mann-Whitney U test. Three-group comparisons with paired donor designs (soft PA, stiff PA, Dynabeads™) were analyzed using the Friedman test with Dunn’s post hoc correction. All analyses were conducted in GraphPad Prism 10 (GraphPad Software, USA). Statistical significance is reported as *p < 0.05, **p < 0.01, ***p < 0.001.

## 3. Results and discussion

### 3.1 Soft and stiff PA platforms decouple stiffness from geometry and ligand density

A central requirement for interrogating mechanical effects is a system in which substrate stiffness can be varied independently of ligand presentation, surface chemistry, and geometry. Polyacrylamide (PA) fulfills this requirement, as its crosslinking density can be tuned across the kPa range relevant to lymphoid tissue while maintaining constant surface chemistry and antibody coupling density [40, 41]. We therefore established a matched-pair PA platform spanning compliant (∼1 kPa) and stiff (∼55 kPa) conditions.

Macroscopic shear rheology **(Figure 1A)** confirmed that these formulations yield mechanically distinct substrates, with steady-state storage moduli of 1.248 ± 0.0108 kPa (soft) and 54.868 ± 1.413 kPa (stiff), and complex viscosities of 0.199 ± 0.00172 kPa·s and 8.732 ± 0.225 kPa·s, respectively. These values span the physiologically relevant compliance of lymphoid tissues through to supra-physiological stiffness, consistent with prior observations of T cell mechanosensitivity [34]. Complementary microscale measurements by AFM **(Figure 1B)** yielded Young’s moduli of 15.05 ± 2.714 kPa (soft) and 49.947 ± 17.776 kPa (stiff). The higher apparent modulus of soft gels by AFM reflects the surface-weighted nature of force spectroscopy relative to bulk rheology. Together, these measurements define the mechanical landscape experienced by T cells at both bulk and interfacial scales [42].

**Figure 1:**
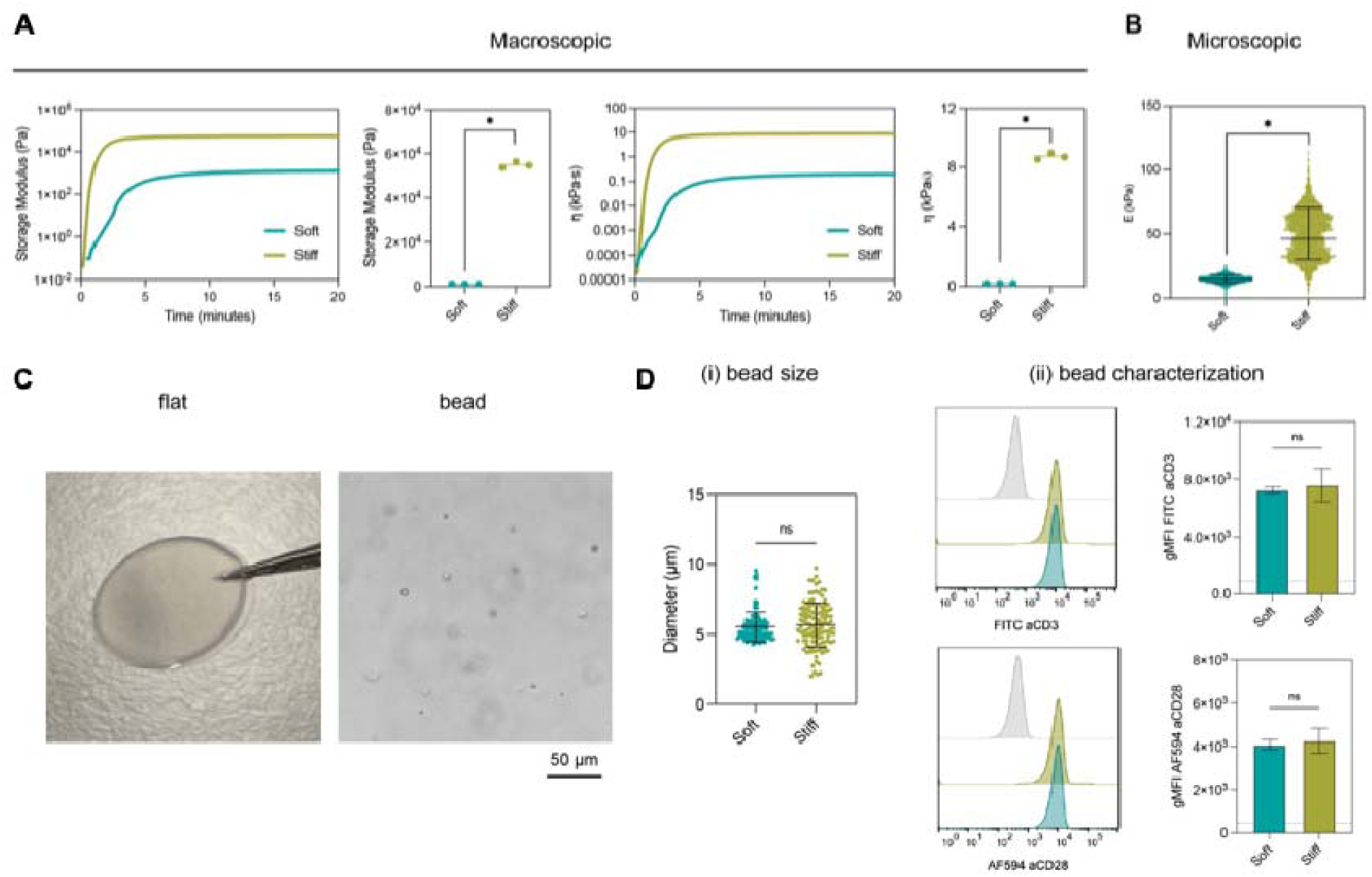
Mechanical and structural characterization of soft and stiff polyacrylamide (PA) platforms. **(A)** Macroscopic viscoelastic properties measured by shear rheology: storage modulus (G′) time-sweep traces and steady-state quantification (left), and complex viscosity (η) time-sweep traces and steady-state quantification (right). **(B)** Microscale Young’s modulus (E) of soft and stiff gels measured by colloidal-probe atomic force microscopy (AFM). **(C)** Representative images of a flat PA gel on a glass coverslip (left) and PA beads in suspension (right); (scale bar=50 µm). **(D) (i)** Diameter distributions of soft and stiff PA beads. (ii) Flow-cytometric histograms (left) and quantified geometric mean fluorescence intensity (gMFI, right) of surface FITC-anti-CD3 and AF594-anti-CD28 on PA beads, demonstrating equivalent antibody functionalization across stiffnesses. Data are mean ± SD of triplicate measurements. Mann-Whitney U test; *p < 0.05; ns, not significant.

To assess whether mechanical effects are preserved across geometries relevant to manufacturing, we generated both flat gels and bead-based formats **(Figure 1C)**. The bead configuration enables evaluation of compatibility with suspension culture systems. Bead diameters were comparable across stiffness conditions (soft 5.55 ± 1.07 µm; stiff 5.68 ± 1.58 µm; **Figure 1D(i))**, within the optimal range for T cell-bead interactions [14]. Importantly, anti-CD3/anti-CD28 coating density was equivalent across soft and stiff beads, as confirmed by flow-cytometric detection of surface-bound antibodies **(Figure 1D(ii))**. Together, these results establish a controlled platform in which stiffness and geometry can be independently varied without confounding differences in ligand density or surface chemistry, providing a robust basis for downstream functional comparisons.

### 3.2 Flat PA outperforms bead geometry at day 7 while Dynabeads™ already show early exhaustion

Whether mechanical input is governed primarily by substrate stiffness or independently shaped by contact geometry remains an open question [18]. The immune synapse (IS) is a spatially organized structure that depends on sustained interfacial contact, with TCR microclusters forming and centralizing over time into mature signalling domains [21, 22]. In this context, bead-based activation provides discontinuous, micron-scale contacts, whereas flat substrates offer a continuous interface across the T cell footprint,a geometric distinction likely to influence signalling quality. To directly test this, we stimulated primary human CD8+ T cells for 7 days on soft and stiff PA substrates in both flat and bead formats, using Dynabead [22]s™ as a clinical-standard comparator.

Flat PA substrates supported compact cluster formation comparable to Dynabeads™, whereas PA beads yielded fewer clusters and significantly lower cell numbers than stiffness-matched flat substrates **(Figure 2A)**, despite identical ligand density. This difference was reflected in activation marker expression **(Figure 2B)**. CD69 induction was similar between Dynabeads™ and flat PA (∼1.4-1.5-fold) but reduced on stiff PA beads (∼0.85-fold). In contrast, CD44 expression was highest on soft flat PA (∼2.3-fold), significantly exceeding both Dynabeads™ and soft PA beads. As CD69 reflects proximal TCR signalling and CD44 marks effector differentiation [43, 44], these results indicate that flat presentation more effectively drives activation and differentiation.

**Figure 2:**
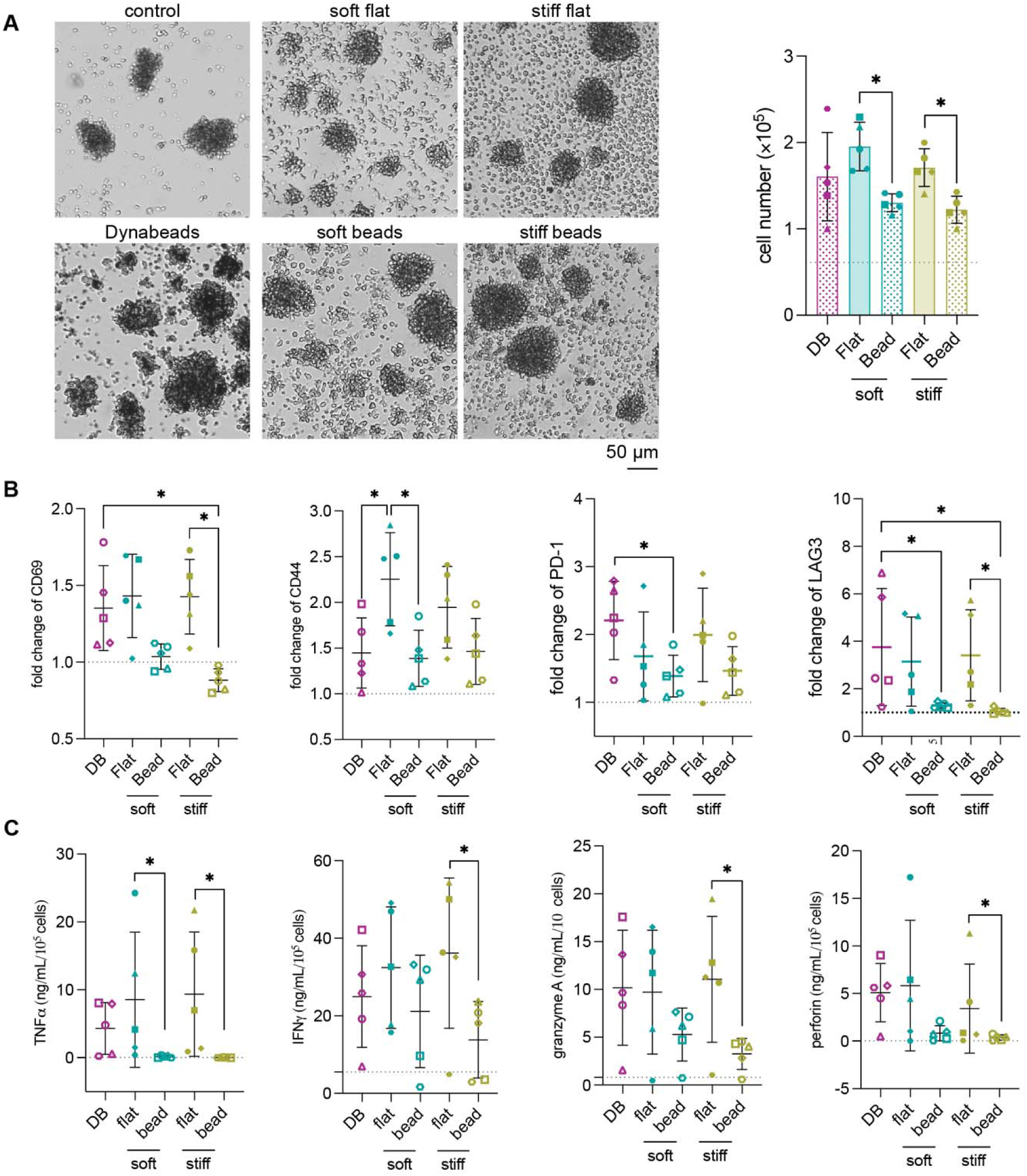
Substrate geometry shapes CD8^+^ T cell activation at day 7. Primary human CD8^+^ T cells were activated for 7 days on Dynabeads™ (DB), soft or stiff flat PA gels, or soft or stiff PA beads, with an unstimulated control. **(A)** Representative brightfield images of cluster formation (scale bar=50 µm) and total cell number at day 7; dotted line, unstimulated control mean. **(B)** Fold-change of CD69, CD44, PD-1, and LAG-3 expression relative to unstimulated control. **(C)** Secretion of TNF-α, IFN-γ, granzyme A, and perforin in culture supernatants. Data are mean ± SD from n = 5 donors with duplicate measurements. Friedman test with Dunn’s post hoc correction; *p < 0.05.

Effector function followed the same geometric trend **(Figure 2C)**. Secretion of TNF-α, IFN-γ, granzyme A, and perforin was significantly higher on stiff flat PA than on stiffness-matched beads, with stiff flat PA achieving the highest absolute levels (IFN-γ 35 ng/mL per 10^5^ cells; granzyme A 11 ng/mL per 10^5^ cells; perforin 3 ng/mL per 10^5^ cells). These molecules represent the core cytotoxic machinery of CD8^+^ T cells [45, 46]. In contrast, Dynabeads™-expanded cells already exhibited elevated exhaustion markers at day 7, with the highest PD-1 (∼2.2-fold) and LAG-3 (∼4-fold) expression across all conditions **(Figure 2B)**, consistent with accelerated checkpoint upregulation under sustained bead stimulation [13, 14].

Mechanistically, these findings suggest that discontinuous bead contacts, irrespective of stiffness, limit full IS maturation, reducing force-dependent TCR engagement and sustained signalling [19]. In contrast, flat PA substrates provide a continuous interface that supports microcluster centralization and stable synapse formation [21, 22]. Although bead-based systems offer practical advantages for suspension culture, planar presentation at physiologically relevant stiffness more effectively drives early activation at matched ligand density. Based on these results, flat PA substrates were prioritized for subsequent analyses. Hereafter, “soft PA” and “stiff PA” refer to the flat configuration unless otherwise specified. Importantly, this comparison also clarifies how distinct physical parameters contribute to T cell activation. Together, these data indicate that contact geometry primarily shapes early activation by influencing immune synapse formation and signaling quality, whereas substrate stiffness becomes the dominant determinant of longer-term T cell behavior in the following sections.

### 3.3 PA substrates sustain balanced activation across 14 days as Dynabeads™ drive PD-1/LAG-3 acquisition and a more pronounced secretory decline

A single time-point analysis is insufficient to determine whether an activation platform sustains T cell function across the full expansion window used in ACT manufacturing. Rapid expansion protocols (REP) typically extend to 14 days, and the phenotype at infusion,rather than the early activation peak,ultimately determines therapeutic efficacy [47]. We therefore profiled CD8^+^ T cell responses to soft PA, stiff PA, and Dynabeads™ longitudinally over 14 days, sampling at days 2, 4, 7, 10, 12 and 14.

We quantified expansion as population doubling level (PDL) **(Figure 3A)** rather than absolute cell count, because PDL is the standard metric for ex vivo T cell expansion in ACT manufacturing, normalizes across donors with different effective proliferative baselines, and maps proliferation onto therapeutically meaningful thresholds, with ∼7-10 PDL considered the optimal therapeutic expansion window and >15 PDL associated with terminal differentiation [10, 48]. Expansion dynamics revealed an early advantage for Dynabeads™. PDL reached ∼6.2 on Dynabeads™ by day 2, compared with a lower range of ∼2.5-3.3 on PA conditions (soft ∼3.3, stiff ∼2.5), and peaked at ∼9 by day 4 on Dynabeads™. In contrast, PA conditions entered the 7-10 PDL range at day 4 (soft ∼8, stiff ∼7), with all conditions converging at ∼10 PDL by day 7. Beyond this point, however, the trajectories diverged. PA-expanded cells maintained ∼10 PDL through day 14, whereas Dynabeads™-expanded cells contracted to ∼6 PDL, indicating net cell loss during the second week. This pattern is consistent with reduced durability rather than excessive proliferation, aligning with clinical observations that terminally differentiated products exhibit poor persistence [10, 12]

**Figure 3:**
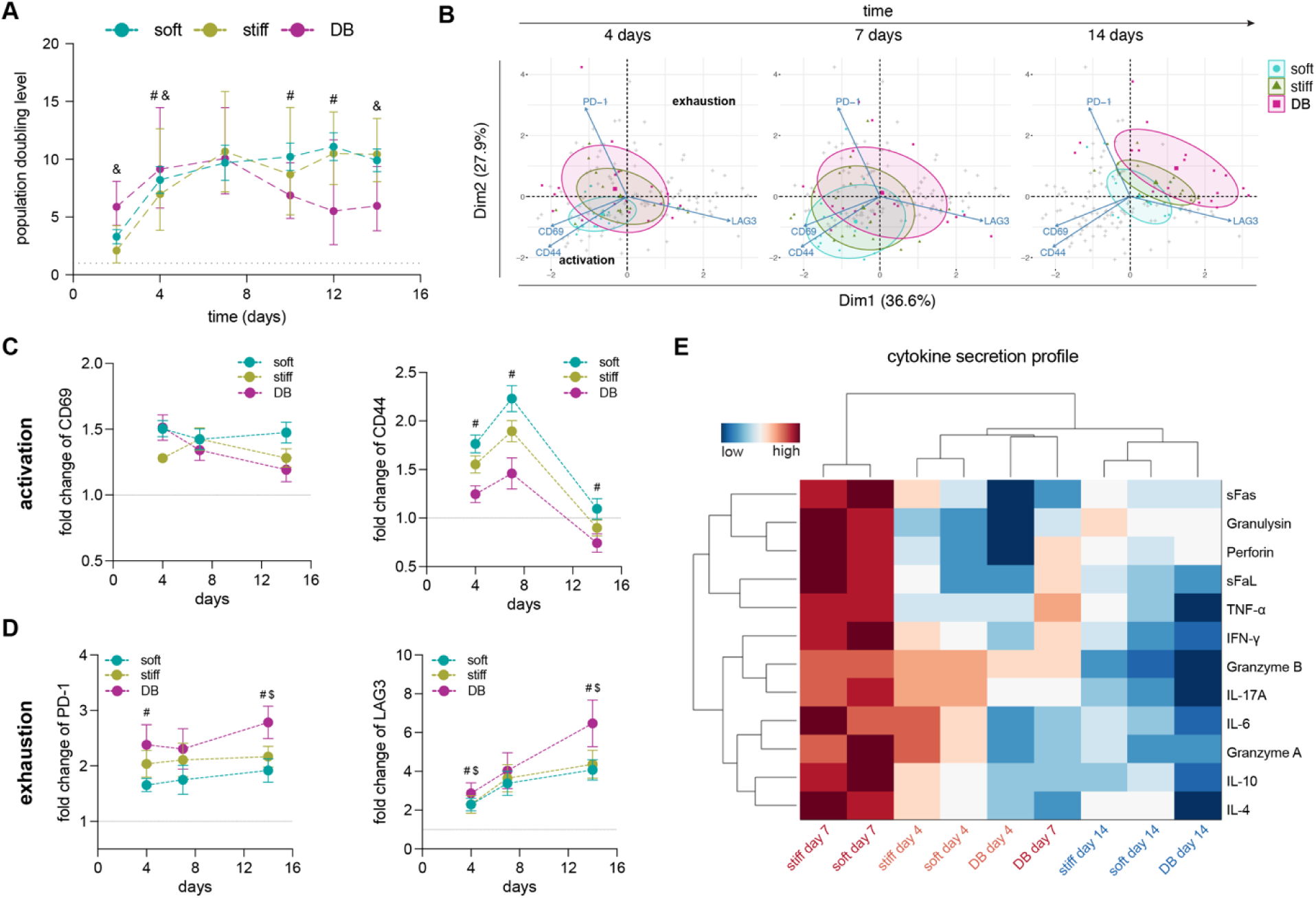
Longitudinal dynamics of CD8^+^ T cell activation, exhaustion, and cytokine secretion over 14 days on soft PA, stiff PA, or Dynabeads™. **(A)** Population doubling level (PDL) at days 2, 4, 7, 10, 12 and 14; dotted line, unstimulated control mean. **(B)** Principal component biplots of CD69, CD44, PD-1, and LAG-3 trajectories at days 4, 7, and 14; ellipses denote 95% confidence intervals for each condition. **(C)** Activation marker kinetics for CD69 (left) and CD44 (right). **(D)** Exhaustion marker kinetics for PD-1 (left) and LAG-3 (right). **(E)** Hierarchical clustergram of 12 cytokine secretion profiles across the nine condition × time-point samples; scale, row-normalized relative abundance. Data are mean ± SD from n = 5 donors with duplicate measurements. Friedman test with Dunn’s post hoc correction. In (A), (D) and (E) (#) soft vs DB; ($) stiff vs DB, *p < 0.05.

To capture multidimensional activation states, we performed principal component analysis of CD69, CD44, PD-1, and LAG-3 trajectories **(Figure 3B)**. At day 4, all conditions occupied overlapping regions, indicating comparable early activation. By day 7, Dynabeads™ shifted along the PD-1/LAG-3 axis, while PA conditions remained closer to the activation axis. By day 14, this separation increased further, with Dynabeads™ progressing toward a checkpoint-dominated state, whereas PA conditions retained a more balanced phenotype. Marker kinetics **(Figure 3C,D)** supported these trends. CD69 declined gradually across all conditions, with soft PA consistently maintaining slightly higher levels. CD44 peaked at day 7 on soft PA (∼2.2), exceeding stiff PA (∼1.9) and Dynabeads™ (∼1.45), before returning toward baseline by day 14. In contrast, PD-1 increased steadily on Dynabeads™ to ∼2.8 by day 14, significantly above soft PA at day 4 and above both PA conditions at day 14 **(Figure 3D)**, while plateauing at ∼2.0 on PA conditions. LAG-3 exhibited the most pronounced divergence, reaching ∼6.5 on Dynabeads™ versus ∼4.0-4.2 on PA, with Dynabeads™ significantly exceeding both PA conditions at days 4 and 14 **(Figure 3D)**. Given that PD-1/LAG-3 co-expression is a hallmark of exhaustion and predicts poor persistence [49, 50], these dynamics indicate accelerated exhaustion under Dynabeads™ stimulation.

Cytokine secretion profiles **(Figure 3E)** further resolved functional trajectories. Hierarchical clustering identified three distinct groups: (i) day-7 PA samples (soft and stiff), representing peak functional output; (ii) intermediate-output samples comprising day-4 conditions and Dynabeads™ day 7; and (iii) low-output day-14 samples across all conditions. At day 4, PA conditions already produced higher levels of TNF-α, IFN-γ, and cytotoxic mediators than Dynabeads™. The day-7 PA peak additionally included elevated perforin, granulysin, IL-4, and IL-10. Notably, IL-4 and IL-10 can enhance CD8^+^ T cell metabolic fitness and resistance to dysfunction [51-53], indicating a balanced effector-regulatory profile rather than suppressed activity. By day 14, cytokine production declined across all conditions; however, PA-expanded cells retained relatively higher IL-4 and IL-10 levels, consistent with a less pronounced functional decline.

Taken together, these longitudinal data reveal a clear divergence in activation trajectories. Dynabeads™ drive rapid early expansion followed by checkpoint accumulation, metabolic decline, and reduced durability, whereas PA substrates sustain activation while limiting exhaustion across the full expansion window. This divergence is most pronounced at day 14, the clinically relevant infusion time point,suggesting that mechanical priming shapes not only early activation but the long-term functional fate of CD8^+^ T cells.

### 3.4 RNA-seq reveals a 125-gene biomimetic core and a Tpex stem-like signature unique to PA-expanded cells

The phenotypic and functional differences observed between platforms suggest a qualitative divergence in underlying transcriptional programs. To test this directly, we performed bulk nanopore RNA-seq on CD8^+^ T cells expanded for 7 days on Dynabeads™, soft PA, or stiff PA, using three independent donors per condition **(Figure 4)**. Day 7 was selected to capture peak functional engagement prior to divergence in late-stage exhaustion trajectories. Principal component analysis **(Figure 4A)** resolved clear separation between conditions along both PC1 (17.6% variance) and PC2 (15.9%). Dynabeads™ samples clustered tightly near the origin, indicating a relatively constrained transcriptional state, whereas PA-expanded cells showed broader dispersion. Soft PA exhibited modest variability, while stiff PA displayed the greatest spread, suggesting increased transcriptional responsiveness in this condition.

**Figure 4:**
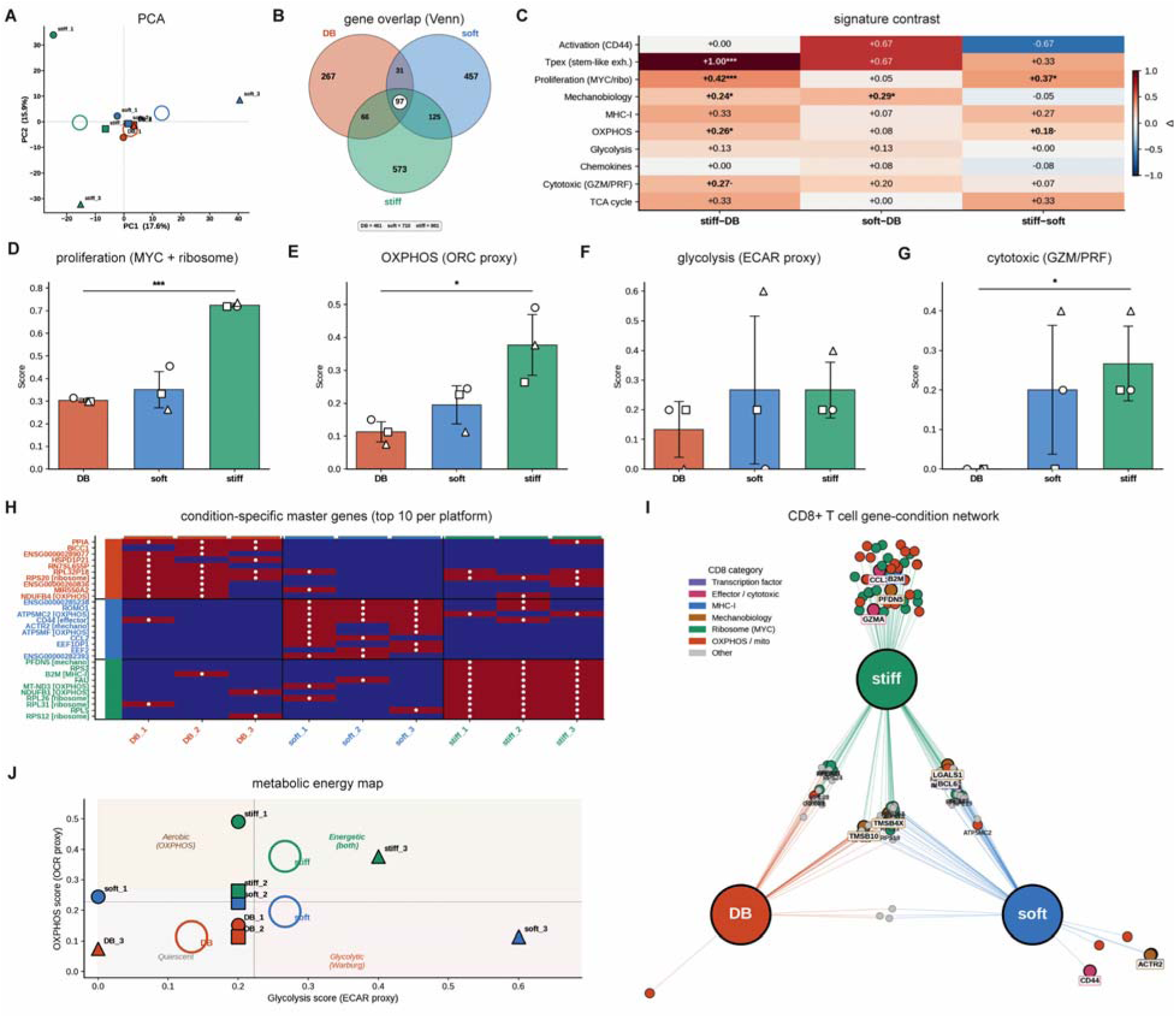
Bulk nanopore RNA-seq identifies a biomimetic transcriptional core shared by soft and stiff PA but absent from Dynabeads™. CD8^+^ T cells expanded for 7 days on Dynabeads™ (DB), soft PA, or stiff PA were profiled in biological triplicate from independent donors. **(A)** Principal component analysis of variance-stabilized gene counts; open circles denote condition centroids. **(B)** Venn diagram of condition-associated gene sets, highlighting the 125-gene “biomimetic core” shared exclusively between soft and stiff PA. **(C)** Pairwise signature contrasts (Δ signature scores) across ten curated gene sets. Statistical significance determined by permutation testing over signature gene membership: ·p < 0.1, *p < 0.05, **p < 0.01, ***p < 0.001. **(D-G)** Per-condition signature scores (mean ± SD across n = 3 donors) for proliferation (MYC + ribosome), OXPHOS (OCR proxy), glycolysis (ECAR proxy), and cytotoxicity (granzyme/perforin); individual donor values overlaid (circle, square, triangle). **(H)** Heatmap of the top 10 condition-specific master genes per platform across the nine samples; red, elevated; blue, depleted; white dots indicate top-ranked drivers. **(I)** Bipartite gene-condition network with nodes colored by functional category. **(J)** Transcriptional metabolic energy map (OXPHOS vs glycolysis signature scores) partitioned into quiescent, aerobic, glycolytic, and energetic quadrants; filled shapes denote individual donors, open circles denote condition centroids.

To define shared and condition-specific programs, we performed gene-set overlap analysis **(Figure 4B)**. This revealed a 125-gene “biomimetic core” shared exclusively between soft and stiff PA but absent from Dynabeads™, exceeding the combined overlap between Dynabeads™ and either PA condition. Only 97 genes were common to all three platforms, representing a minimal shared activation program. Consistent with this, Dynabeads™ exhibited the smallest set of condition-specific genes (267), compared with 457 for soft PA and 573 for stiff PA. These patterns indicate that PA substrates engage a distinct and coordinated transcriptional program, rather than simply amplifying a common response.

Signature-level analysis across ten curated gene sets **(Figure 4C)** further resolved thes differences. Stiff PA showed the strongest enrichment in proliferation (MYC + ribosome; Δ = +0.42 vs DB, p < 0.001), OXPHOS (Δ = +0.26, p < 0.05), and cytotoxicity (Δ = +0.27), whereas soft PA exhibited more moderate increases (Δ = +0.05, +0.08, and +0.20, respectively). Mechanobiology signatures wer comparably enriched in both PA conditions (stiff Δ = +0.24, soft Δ = +0.29; p < 0.05). The most striking contrast was observed in the stem-like precursor (Tpex) signature [54, 55]: stiff PA (Δ = +1.00, p < 0.001) and soft PA (Δ = +0.67) both showed strong enrichment, whereas Dynabeads™ displayed minimal signal. Notably, the CD44 activation signature was higher in soft than in stiff PA, consistent with th phenotypic data in **Figure 3C**.

Per-condition signature scores **(Figure 4D-G)** quantified these trends. Proliferation and OXPHOS increased progressively from Dynabeads™ to soft PA to stiff PA, while glycolysis scores were similar across PA conditions and lower on Dynabeads™. Cytotoxicity signatures were near baseline on Dynabeads™ but elevated on both PA conditions, consistent with reduced effector output observed previously. These data indicate that PA substrates engage both metabolic and functional transcriptional programs that are diminished under bead-based activation.

Master-gene analysis **(Figure 4H,I)** linked these signatures to specific transcriptional drivers. Dynabeads™ were enriched for housekeeping and basal ribosomal/mitochondrial genes, consistent with a less specialized state. Soft PA was characterized by genes associated with cytoskeletal remodeling and effector activation (e.g., CD44, ACTR2, CCL2), whereas stiff PA showed strong enrichment for ribosomal and mitochondrial genes (e.g., RPL26, RPS3, MT-ND3, NDUFB1), indicative of elevated biosynthetic and metabolic activity. Several genes (TMSB4X, TMSB10, LGALS1, BCL6) were shared between PA conditions but absent from Dynabeads™, forming key components of the biomimetic core.

Finally, a transcriptional metabolic map **(Figure 4J)** positioned Dynabeads™ in a quiescent state (low OXPHOS, low glycolysis), soft PA at the interface between aerobic and glycolytic states, and stiff PA in a highly energetic, dual-engaged state. As these signatures serve as transcriptional proxies for OCR and ECAR, respectively, this mapping provides a functional prediction for metabolic behavior across the expansion timeline.

Together, these data identify a biomimetic transcriptional program uniquely engaged by PA substrates, characterized by coordinated activation, metabolic engagement, and enrichment of a Tpex stem-like state associated with durable T cell responses [54, 55].

### 3.5 Soft PA builds day-14 bioenergetic flexibility while Dynabeads™ show the lowest metabolic output

The transcriptional metabolic map **(Figure 4J)** predicted reduced metabolic engagement in Dynabeads™-expanded cells and differential activation of oxidative and glycolytic programs across PA conditions. To test these predictions and define their temporal evolution, we performed Seahorse extracellular flux analysis across the 14-day expansion window **(Figure 5)**. As T cell effector function is tightly coupled to metabolic state, relying on glycolysis for biosynthesis and oxidative phosphorylation (OXPHOS) for persistence, this analysis provides a functional readout of metabolic fitness [56-58] **(Figure 5A)**

**Figure 5:**
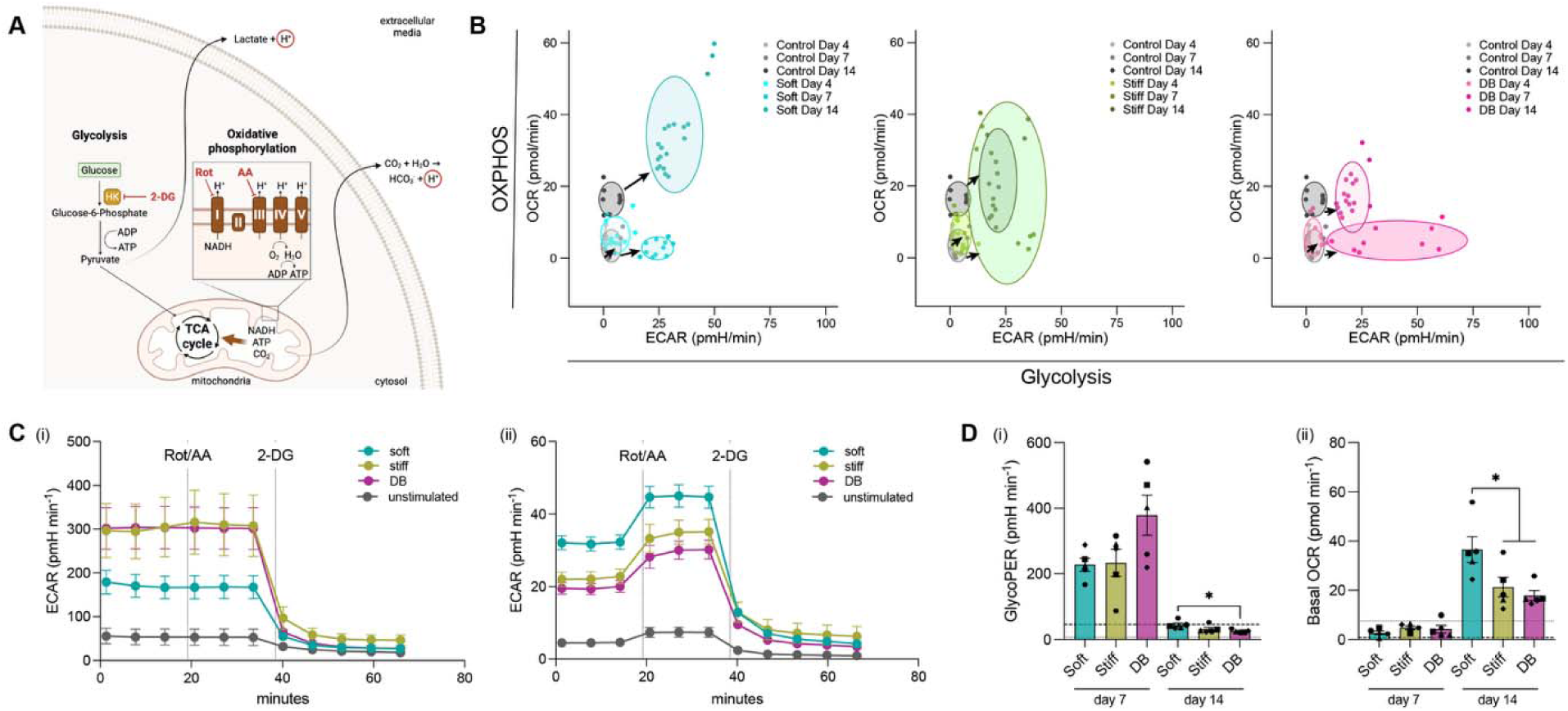
Seahorse extracellular flux analysis reveals divergent metabolic trajectories across 14 days. CD8^+^ T cells expanded on soft PA, stiff PA, or Dynabeads™ were profiled at days 4, 7, and 14. **(A)** Schematic of glycolysis and oxidative phosphorylation pathways targeted by sequential injection of rotenone/antimycin A (Rot/AA) and 2-deoxy-D-glucose (2-DG). **(B)** OCR-versus-ECAR energy diagrams at days 4, 7, and 14 for soft PA (left), stiff PA (center), and Dynabeads™ (right); arrows indicate trajectories relative to the unstimulated control centroid. **(C)** ECAR traces during sequential Rot/AA and 2-DG injection at day 7 **(i)** and day 14 **(ii)**. (D) Quantified glycolytic proton efflux rate (GlycoPER) **(i)** and basal OCR **(ii)** at days 7 and 14. Data are mean ± SD from n = 5 donors with triplicate measurements. Friedman test with Dunn’s post hoc correction; *p < 0.05.

At day 4, all conditions clustered near the origin of the OCR-ECAR energy map **(Figure 5B)**, consistent with early metabolic priming prior to full effector commitment. By day 7, the platforms diverged. Dynabeads™ and stiff PA shifted toward a glycolytic phenotype, with Dynabeads™ and stiff PA reaching comparably high ECAR (∼300 mpH/min each) and soft PA the lowest (∼175 mpH/min**) (Figure 5C(i))**. Basal OCR remained low across conditions **(Figure 5D(ii)**, left), indicating that transcriptional engagement of OXPHOS had not yet translated into increased respiratory output. This lag suggests a temporal decoupling between transcriptional activation and metabolic execution. By day 14, the metabolic states had diverged more substantially. Soft PA-expanded cells occupied the energetic quadrant, exhibiting the highest basal OCR (∼35 pmol/min) together with sustained ECAR (∼40 mpH/min) **(Figure 5C(ii), D(ii))**. This dual engagement reflects a state of bioenergetic flexibility, a metabolic phenotype previously associated with CD8^+^ memory T cell formation and long-term survival [58]. Stiff PA maintained intermediate activity (OCR ∼20 pmol/min; ECAR ∼30 mpH/min), whereas Dynabeads™-expanded cells exhibited the lowest values on both axes (OCR ∼15 pmol/min; ECAR ∼20 mpH/min), consistent with reduced metabolic capacity. Although GlycoPER declined across all conditions from day 7 to day 14, inter-condition differences at day 14 were driven primarily by mitochondrial respiration, with soft PA showing the strongest OXPHOS engagement.

The Dynabeads™ metabolic profile,low OCR and low ECAR at day 14,is characteristic of terminal metabolic exhaustion. Mechanistically, sustained PD-1 signalling suppresses glycolysis and impairs mitochondrial biogenesis, leading to energetically inefficient states [5, 59]. In contrast, soft PA-expanded cells retain engagement of both glycolytic and oxidative pathways, consistent with metabolic states associated with durable memory and persistence [58, 60]. Clinical evidence further supports the relevance of this phenotype, linking metabolic fitness at infusion to improved CAR-T cell persistence and tumor control [61-63]. Notably, the transcriptional and functional data exhibit a temporal offset. At day 7, OXPHOS-associated transcriptional programs are most enriched in stiff PA **(Figure 4E)**, whereas by day 14 soft PA exhibits the highest respiratory output. This divergence is consistent with delayed metabolic realization on compliant substrates, although direct measurements of mitochondrial mass or spare respiratory capacity would be required to confirm this mechanism. Together, these data define distinct metabolic trajectories across platforms. Dynabeads™ induce early glycolytic activation followed by metabolic decline, stiff PA supports sustained intermediate activity, and soft PA progressively establishes a balanced, dual-pathway metabolic state that peaks at day 14. This bioenergetic flexibility aligns with the clinically relevant infusion time point, linking mechanical priming to metabolic fitness and, by extension, durable anti-tumor function.

### 3.6 PA priming sustains CD8^+^ T cell cytotoxicity against MDA-MB-231 and MCF-7 in 2D and under 3D collagen constraints

The ultimate test of mechanical priming is whether the activation, transcriptional, and metabolic advantages of PA substrates translate into durable cytotoxic function. We therefore evaluated CD8^+^ T cells expanded for 4, 7, or 14 days against MDA-MB-231 (triple-negative) and MCF-7 (luminal) breast cancer cells in both 2D and collagen-based 3D co-culture systems. Imaging confirmed effective target engagement across conditions **(Figures 6A, 7A)**. In 2D, PA-expanded T cells formed compact clusters and disrupted tumor monolayers, while in 3D they infiltrated collagen matrices and formed immune synapses characterized by localized F-actin accumulation at the T cell-tumor interface,a hallmark of productive cytotoxic engagement [24, 64].

**Figure 6:**
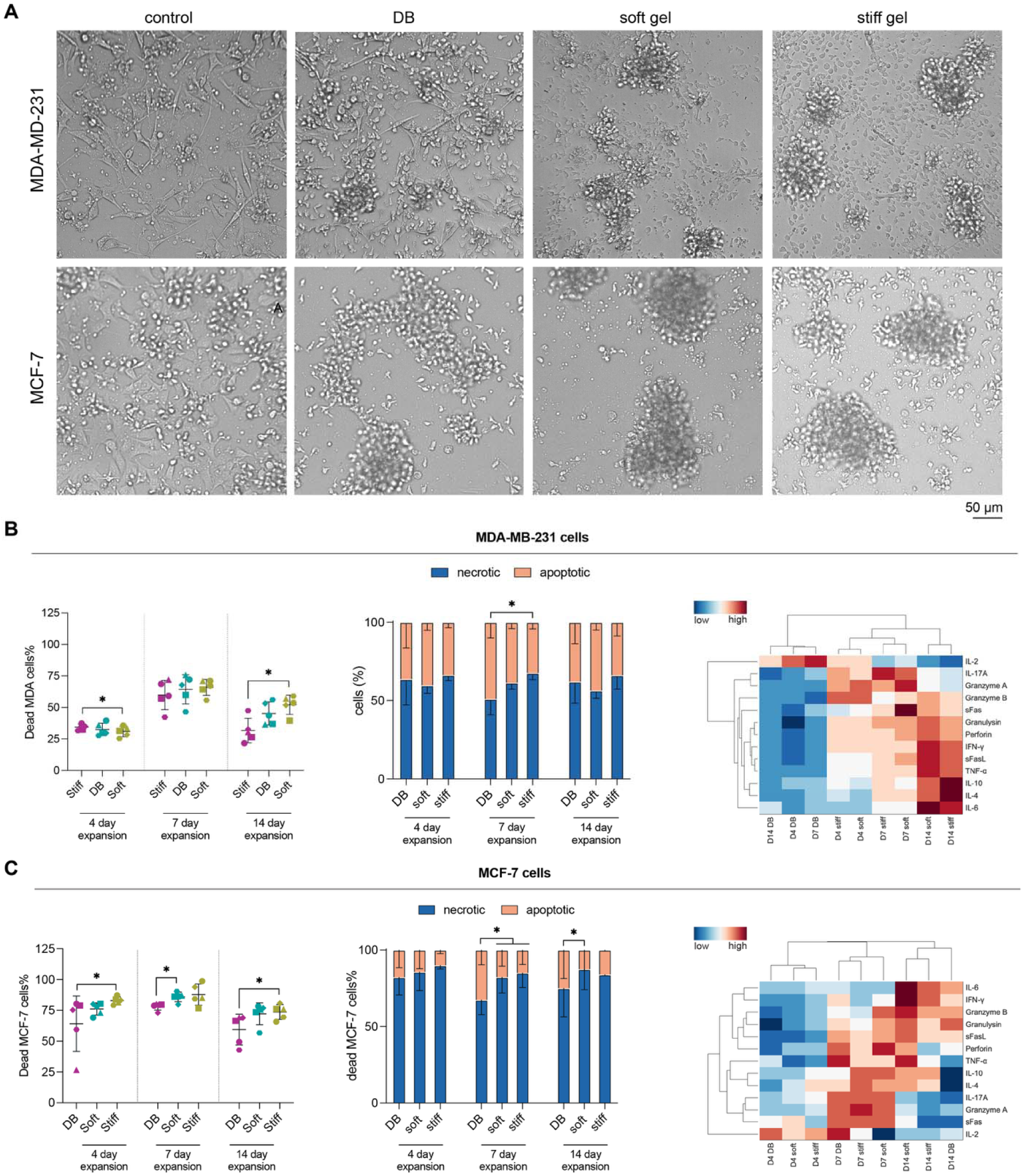
CD8^+^ T cell-mediated killing of MDA-MB-231 and MCF-7 breast cancer cells in 2D co-culture. CD8^+^ T cells expanded on soft PA, stiff PA, or Dynabeads™ (DB) for 4, 7, or 14 days were co-cultured with tumor cells for 72 h at a 3:1 effector-to-target ratio. **(A)** Representative brightfield images of tumor cell disruption by day-14-expanded T cells against MDA-MB-231 (top row) and MCF-7 (bottom row); control = tumor cells without T cells. (Scale bar=50 µm). **(B)** MDA-MB-231 co-culture outcomes: **(i)** percentage of dead tumor cells across expansion time points; **(ii)** apoptotic (caspase-3/7^+^DRAQ7^-^) versus necrotic (DRAQ7^+^) fractions; **(iii)** hierarchical clustergram of secreted cytokine profiles across the nine condition × time-point samples. **(C)** MCF-7 co-culture outcomes: matched panels. Data are mean ± SD from n = 5 donors with duplicate measurements. Friedman test with Dunn’s post hoc correction; *p < 0.05.

In 2D co-culture **(Figure 6B,C)**, all conditions exhibited comparable killing at day 7, representing peak cytotoxic output. However, clear divergence emerged at day 14. Against MDA-MB-231, soft PA retained ∼52% killing and Dynabeads™ retained ∼45%, whereas stiff PA significantly declined to ∼30%. Against MCF-7, overall killing was higher, peaking at ∼85% on day 7 across conditions; by day 14, PA conditions retained ∼70-75% killing compared with a more pronounced decline to ∼60% for Dynabeads™. These results indicate that the primary advantage of PA priming lies not in increasing peak cytotoxicity, but in preserving function over time. In 3D collagen co-culture **(Figure 7B,C)**, overall killing was reduced across all conditions, consistent with matrix-imposed constraints on migration and synapse formation. Despite this, the PA advantage at day 14 was preserved. Against MDA-MB-231, killing was similar across conditions at days 4 and 7 (day 4 ∼40%; day 7 ∼50%) but diverged at day 14, with soft and stiff PA both retaining ∼45% killing compared with ∼37% for Dynabeads™. Against MCF-7, PA conditions maintained ∼80% killing across time points, whereas Dynabeads™ declined to ∼60% at day 14. Thus, mechanical priming confers a durability advantage even under conditions that more closely mimic the solid tumor microenvironment.

**Figure 7:**
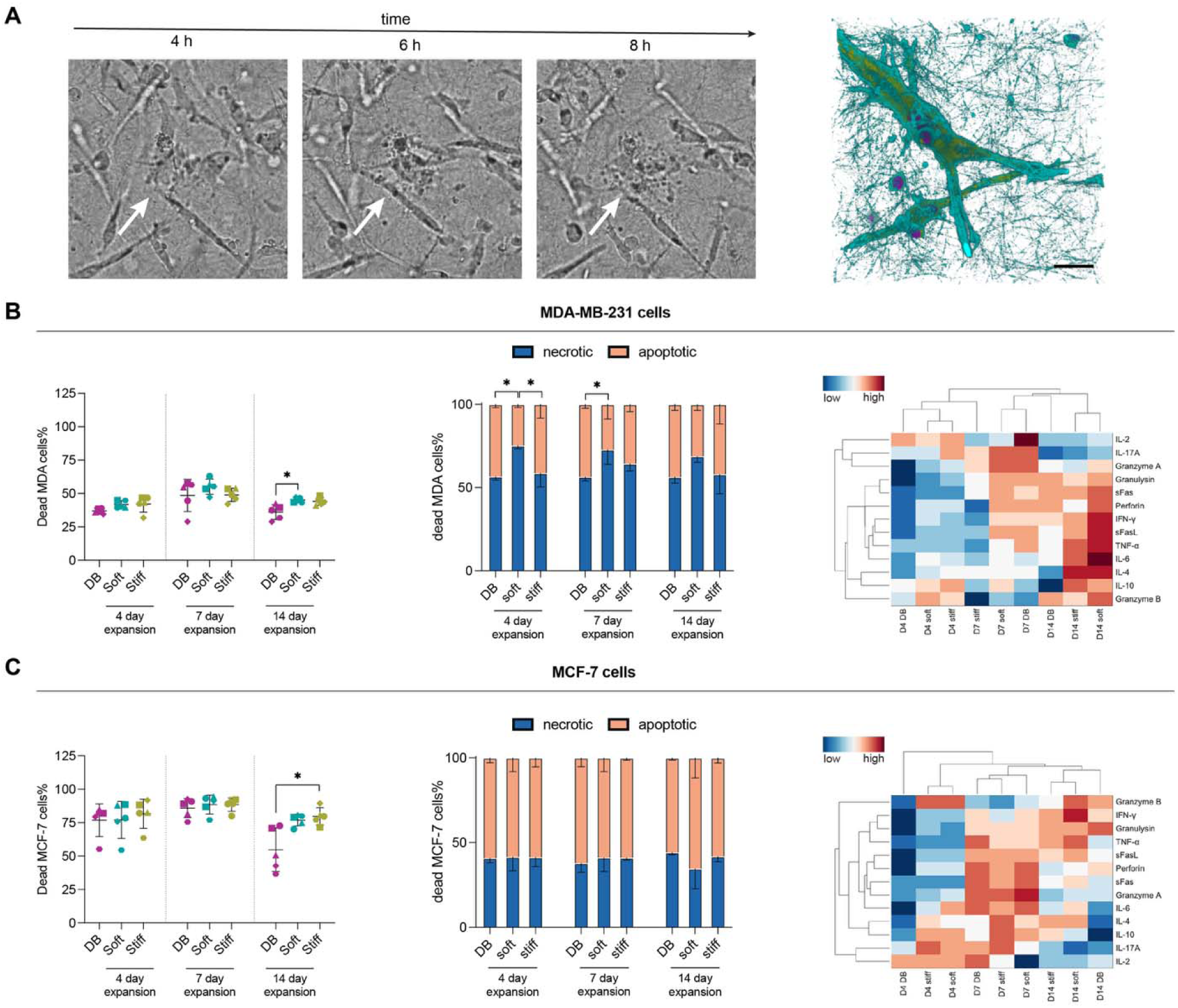
CD8^+^ T cell-mediated killing of MDA-MB-231 and MCF-7 breast cancer cells in 3D collagen co-culture. CD8^+^ T cells expanded on soft PA, stiff PA, or Dynabeads™ (DB) for 4, 7, or 14 days were co-cultured with tumor cells embedded in 2 mg/mL reconstituted collagen type I matrices for 72 h at a 3:1 effector-to-target ratio. **(A)** Left: time-lapse brightfield images (4 h, 6 h, 8 h) showing a CD8^+^ T cell (arrow) engaging and disrupting an MDA-MB-231 cell within the collagen matrix. Right: confocal 3D reconstruction of a cytotoxic immune synapse with localized F-actin enrichment at the T cell-tumor interface (Hoechst nuclei, magenta; phalloidin F-actin, cyan; α-tubulin, yellow). (Scale bar=10 µm). **(B)** MDA-MB-231 3D co-culture outcomes: **(i)** percentage of dead tumor cells across expansion time points; **(ii)** apoptotic versus necrotic fractions; **(iii)** hierarchical clustergram of secreted cytokine profiles. (C) MCF-7 3D co-culture outcomes: matched panels. Data are mean ± SD from n = 5 donors with duplicate measurements. Friedman test with Dunn’s post hoc correction; *p < 0.05.

Analysis of killing modality provided further insight. In MDA-MB-231 3D co-culture, PA-expanded cells exhibited a higher proportion of apoptotic killing at early time points, whereas Dynabeads™-expanded cells showed relatively greater necrotic fractions **(Figure 7B)**. Apoptotic killing reflects coordinated perforin/granzyme-mediated cytolysis, while necrotic killing is associated with impaired synapse function or inefficient granule delivery [45, 46, 65]. These differences diminished by day 14, suggesting that modality shifts are most pronounced during early functional engagement. Cytokine profiling of co-cultures further supported these trends. Across both 2D and 3D systems, PA-expanded cells exhibited higher effector cytokine output at day 14, whereas Dynabeads™ samples consistently clustered with the lowest-output group. Notably, PA conditions retained elevated IL-4 and IL-10 at day 14, cytokines associated with sustained CD8^+^ T cell function and resistance to exhaustion [51-53].

Taken together, these results demonstrate that PA priming preserves cytotoxic function over time and under physiologically relevant constraints. While peak killing at day 7 is similar across platforms, PA-expanded T cells maintain this function at day 14, whereas Dynabeads™-expanded cells decline. This durability advantage is conserved across tumor subtypes and persists in matrix-constrained 3D environments, directly linking mechanical programming during expansion to sustained anti-tumor efficacy.

## 4. General Discussion and Conclusion

This study identifies substrate mechanics as a primary determinant of CD8^+^ T cell fate during ex vivo expansion. By isolating stiffness and geometry from biochemical inputs, we show that biomimetic polyacrylamide (PA) substrates program T cells toward a transcriptionally coordinated and metabolically flexible state that sustains cytotoxic function in matrix-constrained tumor models. In contrast, Dynabeads™, the current clinical standard, drive rapid activation followed by checkpoint accumulation, metabolic decline, and functional loss [13, 14]. Across all analytical layers, a consistent pattern emerges. PA-expanded cells preserve a stem-like precursor (Tpex) transcriptional program, maintain balanced effector-regulatory cytokine output, and develop bioenergetic flexibility characterized by sustained engagement of both glycolytic and oxidative pathways. These features converge functionally in the retention of cytotoxic activity at day 14, particularly under 3D collagen constraints that more closely reflect solid tumor microenvironments [7, 8].

Notably, the advantage of PA substrates lies not in enhancing peak activation, but in preserving functional capacity over time, an outcome directly aligned with clinical requirements for durable ACT responses [10, 12]. Soft and stiff PA substrates exhibit complementary effects. Stiff PA enhances early transcriptional and metabolic engagement, whereas soft PA supports maximal CD44 expression and establishes the most favorable day-14 metabolic state. At day 14, biomimetic PA substrates significantly exceeded Dynabeads™ in three of the four tumor-killing contexts tested: stiff PA against MCF-7 in both 2D and 3D co-cultures, and soft PA against MDA-MB-231 in 3D collagen. In MDA-MB-231 2D, neither PA condition reached a significant difference from Dynabeads™, although soft PA trended higher while stiff PA trended lower, yielding a significant within-PA difference instead. These patterns indicate that biomimetic compliance, rather than a single optimal stiffness, underlies the durable cytotoxic advantage, with soft (∼1 kPa) substrates providing the most context-independent benefit. Notably, the soft substrate condition (∼1 kPa) lies within the 0.2-1.5 kPa range reported for human monocyte-derived dendritic cells [66, 67], raising the possibility that such compliant interfaces better approximate aspects of T cell-APC mechanical coupling compared to rigid bead-based systems. Together, these findings support a model in which substrate compliance tunes the balance between early activation intensity and long-term functional stability.

From a translational perspective, these results highlight a largely unexploited design parameter in ACT manufacturing. Current expansion protocols rely on rigid polystyrene beads, whose mechanical properties are dictated by material constraints rather than biological relevance [30, 31]. Our data suggest that this choice contributes to the development of exhausted, metabolically compromised T cell products. In contrast, mechanical priming via compliant substrates achieves phenotypes associated with clinical efficacy, including reduced PD-1/LAG-3 co-expression, preservation of Tpex states, and sustained oxidative metabolism [50, 58],without requiring modification of cytokine regimens or genetic engineering strategies. As such, substrate mechanics could be integrated into existing GMP workflows as a modular and scalable intervention.

Several limitations warrant consideration. The RNA-seq dataset (n = 3 donors per condition) supports robust signature-level conclusions but limits gene-level resolution. The tumor models used here capture key biological differences but do not encompass the full heterogeneity of solid tumors; in vivo validation in syngeneic or humanized systems will be required to confirm translational relevance. In addition, scaling compliant substrates to manufacturing settings presents engineering challenges, although emerging approaches, including soft-bead systems, perfusion bioreactors, and microfluidic platforms, offer viable pathways forward. Finally, while this study isolates stiffness and geometry, other material properties such as ligand mobility, topography, and viscoelasticity may further refine T cell programming and merit systematic investigation [29].

In summary, we demonstrate that the mechanical environment of the activation platform is not a passive feature of T cell culture, but a programmable input that shapes long-term functional fate. By linking substrate mechanics to transcriptional state, metabolic fitness, and sustained cytotoxicity, this work establishes mechanical priming as a practical and actionable strategy for improving the durability of CD8^+^ T cell therapies in solid tumors.

## Acknowledgments

The authors acknowledge support from the New York University Abu Dhabi (NYUAD) Faculty Research Fund (AD266) and the New York University Abu Dhabi Global Ph.D. Fellowship. The authors would also like to acknowledge support from the NYUAD research consumable stores and NYUAD core technology platform: Light Microscopy and Molecular and Cell Biology.

## CRediT authorship contribution statement

**Aseel Alatoom:** Writing – review & editing, Writing – original draft, Investigation, Formal analysis, Methodology, Data curation, Visualization, Conceptualization. **Jiranuwat Sapudom:** Writing – review & editing, Writing – original draft, Investigation, Formal analysis, Methodology, Supervision, Conceptualization. **Kamal Elkhoury:** Investigation, Formal analysis. **Muhammedin Deliorman:** Investigation, Formal analysis. **Mostafa Khair:** Investigation, Formal analysis. **Sanjairaj Vijayavenkataraman:** Writing – review & editing, Resources. **Mohammad Qasaimeh:** Writing – review & editing, Resources. **Jeremy Teo:** Writing – review & editing, Conceptualization, Supervision, Project administration, Funding acquisition.

## Data Availability

Data are available from the corresponding author upon reasonable request.

